# Supervised restricted data fusion with common, local & distinct components

**DOI:** 10.64898/2026.04.30.721639

**Authors:** Fred T.G. White, Geert Roelof van der Ploeg, Anna Heintz-Buschart, Lemeng Dong, Harro J. Bouwmeester, Age K. Smilde, Johan A. Westerhuis

## Abstract

In multi-block data, the dominant sources of variation are not always most relevant to a response of interest, meaning that purely exploratory decompositions may fail to recover subtle but important response-associated structure. We introduce PESCAR, a supervised extension of *Penalised Exponential Simultaneous Component Analysis* (PESCA) that incorporates response information directly into the estimation of common, local, and distinct (CLD) structure across multiple data blocks. This allows simultaneous multiblock decomposition and response variable influenced recovery of latent structure. Through simulation studies, we show that PESCAR can detect weak response-related components across a range of settings, including different noise levels and model-rank mis-specification. Applied to a real multi-omics dataset, PESCAR recovers biologically meaningful response-associated patterns and retains interpretable block structure. We further demonstrate that sparsity in the fitted loading matrices admits a hypergraph-based interpretability layer, summarising overlapping support patterns across components and blocks. These results show that direct incorporation of response information into multiblock decomposition can improve detection of subtle relevant signal and facilitate interpretation in complex systems.

## Introduction

Analysing multiple high-dimensional data blocks measured on the same samples is now routine in systems biology (1),(2) and psychological science (3),(4). Multi-omics studies, for example, combine transcriptomics, metabolomics, proteomics, and microbiome profiles to interrogate the same biological system; similarly, multimodal studies in psychology and neuroscience integrate *magnetic resonance imaging* (MRI), and *electroencephalography* (EEG), and behavioural assessments. These multiblock datasets provide a more holistic view of the systems under investigation. However, the analysis of these kinds of data is not devoid of challenges in data fusion and model interpretability.

One key aspect in multiblock analysis is the identification of common, local and distinct (CLD) structure across different blocks (5),(6). These structures are essential for understanding the high-level topology of the dataset i.e. the higher-order pattern of overlap and separation among feature sets and data blocks induced by the latent components.

There already exist several data fusion methods including *joint and individual variation explained* (JIVE (7)), *distinct and common simultaneous component analysis* (DISCO (8)), *structural learning and integrative decomposition of multi-view data* (SLIDE(9)), *n-way orthogonal partial least squares* (OnPLS (10), (11)), and *multi-omics factor analysis* (MOFA (12),(13)). These methods differ substantially in formulation, but often face one or more limitations relevant to CLD analysis, including restricted handling of local structure, reliance on rotations or stepwise searches, extensive structure-selection procedures, or difficulties scaling model selection as the number of blocks increases. *Penalised Exponential Simultaneous Component Analysis* (PESCA (6)) provides an attractive alternative because it estimates CLD structure simultaneously through penalised sparse loading matrices, avoiding explicit combinatorial searches over block-sharing patterns. Its computational complexity scales with the number of samples and features rather than with the number of blocks, making it well suited to high-dimensional multiblock settings. In addition, PESCA is formulated within the exponential-family framework and can therefore accommodate not only Gaussian data, but also non-Gaussian data types such as Bernoulli and Poisson observations.

While PESCA and related exploratory methods focus on reconstructing systematic variation, supervised settings require a different emphasis. In regression problems, it is not necessary to describe all systematic variation in the predictors; rather, the goal is to recover the part that is most relevant to a response. In noisy and high-dimensional multiblock data, response-associated structure may be small relative to dominant unsupervised sources of variation, and may therefore be poorly represented by a purely exploratory decomposition.

This creates the need for a method that estimates CLD structure while also describing a response. In particular, such a method should integrate multiblock data in relation to a response or covariate while recovering smaller response-related components in noisy and complex settings.

Existing methods such as *supervised integrated factor analysis* (SIFA (14)), *supervised JIVE* (sJIVE (15)), and *sparse common and distinctive covariate regression* (SCD-CovR (16)) have facilitated supervised data fusion, or at least data fusion in the context of a covariate. sJIVE jointly estimates joint (common) and block-specific (distinct) structure while predicting the outcome, but its decomposition does not account for local variation patterns. In addition, sJIVE requires selection of multiple ranks (one joint rank and a block-specific individual rank for each data block), which becomes increasingly complex as the number of blocks increases. SCD-CovR can represent common, local and distinct components through zero-block constraints where selected block specific parts of the component weight matrix are explicitly fixed to zero; however, this means that the component–block structure is effectively specified and selected through a constrained model search. In practice, multiple influential parameters must be tuned, including the prediction–reconstruction weighting, ridge/lasso penalties, the number of components, and the zero-block constraint configuration, which the authors address via sequential cross-validation because joint tuning over all combinations is computationally costly. This model-selection issue becomes more pronounced as the number of blocks increases and the space of candidate CLD structures expands.

These limitations highlight the need for an alternative approach that preserves the simultaneous estimation strategy of PESCA while incorporating a response variable directly into the model, allowing the recovered CLD structure to be shaped by response-relevant variation. In this work we introduce Penalised Simultaneous Component Analysis Regression (PESCAR) as an answer to this. PESCAR can be viewed from two complementary perspectives: as regression on a response under a CLD-structured penalty, or as a multiblock decomposition in which the recovered CLD structure is guided by variation relevant to a response.

In addition to the supervised extension from PESCA to PESCAR, the fitted model also supports an interpretability layer through hypergraph visualisation (**Fig. 1**). Sparsity in the loading matrices induces a discrete support structure pattern describing which features are active (nonzero) or inactive (zero) in each component. This can be summarised topologically: nodes represent block-specific feature groups, and hyperedges represent shared component-level support patterns. Unlike ordinary edges, which connect pairs of nodes, hyperedges can connect multiple nodes simultaneously; in the visualisation, these are shown as enclosing regions. A hyperedge may contain multiple nodes from the same block, reflecting that a single CLD component can involve several distinct feature groups from that block. Nodes appearing in only one hyperedge indicate component-specific feature groups, whereas nodes appearing in multiple hyperedges indicate feature groups that link components through overlapping support. This representation encourages a systems-level interpretation that is closer to component-based SEM and path-modelling frameworks ((17),(18),(19)), although here it remains a descriptive summary of overlapping support rather than an explicitly directed model.

**Figure 1.**
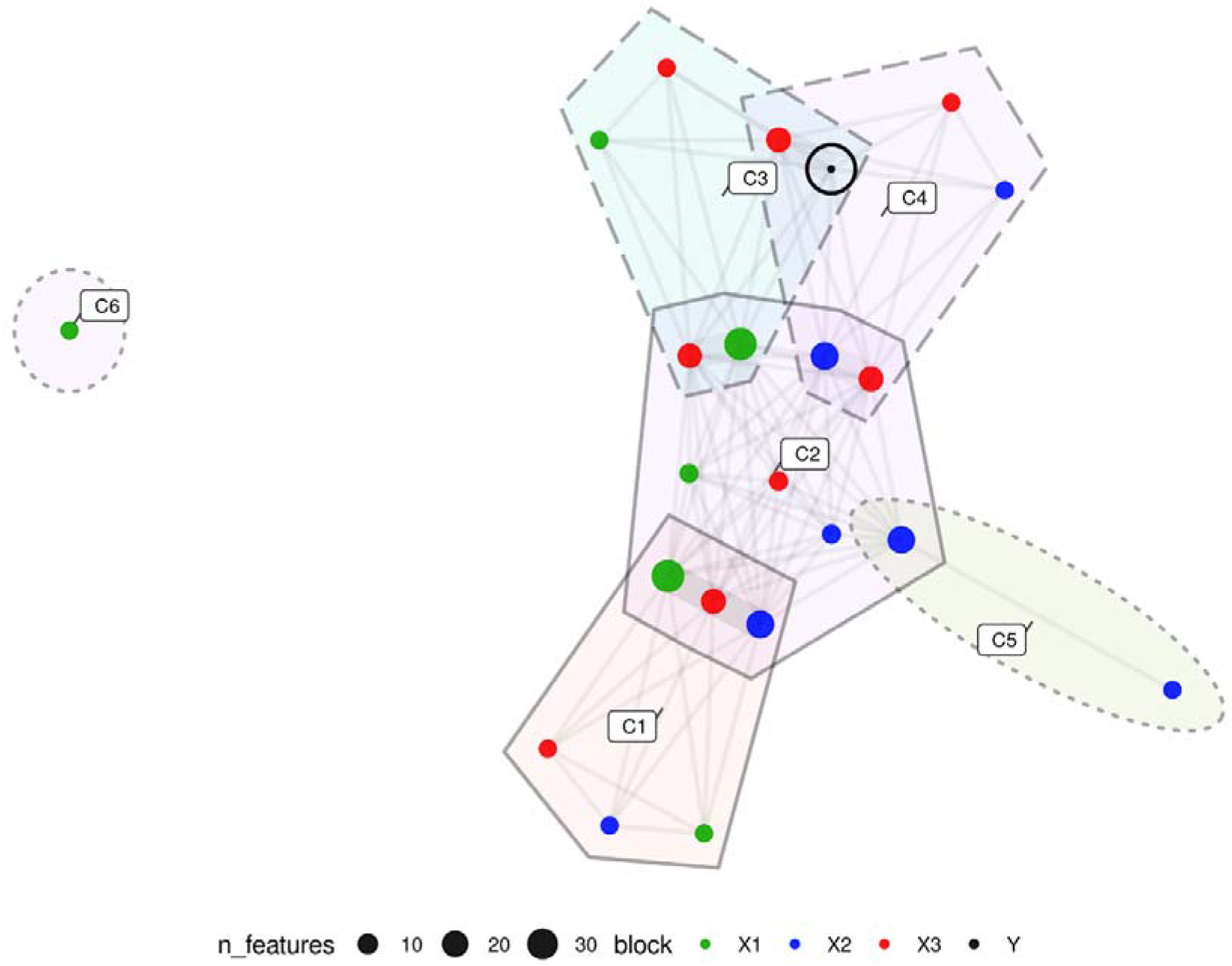
Topological summary of exemplary common, local, and distinct (CLD) components. Shaded regions indicate latent CLD components (hyperedges): each region corresponds to one component classified as Common (C), Local (L), or Distinct (D) according to the set of data blocks represented in that component (C spans all blocks, L spans a subset, D comes from a single block; these types are indicated by solid, long-dashed and short-dashed borders respectively). Nodes represent subsets of features, grouped by block and by their component-membership pattern, Y node is circled in black. An undirected edge between two nodes indicates that the corresponding feature groups co-occur in at least one component; edge width is proportional to the number of components that are nonzero across the connected nodes.

In the following, we describe the mathematics of the proposed model, as well as the algorithm used to estimate the model parameters. We demonstrate and test the approach on simulated datasets and finally demonstrate its applicability on a real multi-omics dataset investigating the relationships between the root transcriptome, metabolome and microbiome in control and nitrogen deficient tomato (20).

## Materials and Methods

### Models

#### PESCA – Penalised Exponential Simultaneous Component Analysis (6)

The standard PESCA model seeks to fit up to L blocks of data **X**_*l*_ (*l* = 1 … *L*) of dimensions I × J_l_ where I is the number of samples and (J_l_) the number of features per block. These blocks are modelled in R components in the standard matrix factorisation as:

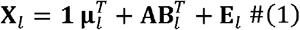

Where the orthogonal scores **A**(I × R) and loadings **B**(J_l_ × R) represent the variation patterns an d the association of each feature to those patterns respectively, with block residual terms in **E**_*l*_.

To estimate this, the loss is formulated as a (multiblock) SCA model which is estimated here by minimising the residual sum of squares 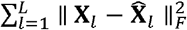 where 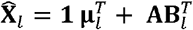 is the rank R model estimation of the data. To induce the block structured sparsity patterns and therefore infer CLD structure, a Generalised Double Pareto (GDP) penalty (21); 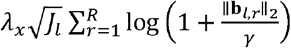 is imposed, where **b**_*l,r*_ is the r^th^ column of **B**_*l*_, *λ*_*x*_ and *γ* are tuning parameters, *λ*_*x*_ controls the overall strength of the penalty and *γ* controls the scale at which the penalty relaxes, smaller *γ* values penalize weak groups (those with low ‖ **b**_*l,r*_‖_2_) more aggressively. *J*_*l*_ enables block specific scaling by the square root of the number of features. Because the penalty acts on ‖ **b**_*l,r*_‖_2_, sparsity occurs at the component block level: if ‖ **b**_*l,r*_‖_2_ = 0, then component *r* contributes no signal to block *l*. The resulting pattern of nonzero columns across blocks directly encodes CLD structure, and this across block sparsity provides an estimate of the effective rank per block. This then gives the full loss function of PESCA as:

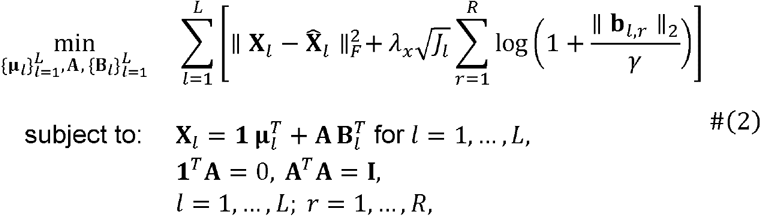

For details loss function gradient the reader is referred to the supplementary materials (section 1) and for more in depth discussion on PESCA model formulation the reader is referred back to (6).

#### PESCAR – PESCA-Regression

In PESCAR simultaneous fitting of L data blocks **X**_*l*_ = [**X**_1_, …, **X**_L_]) and a response (I× 1) is desired. The (centered) model logically becomes:

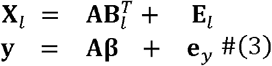

The response **y** is modelled by (a subset of) the same scores **A** as **X** and their respective loadings **B**_*l*_ = [**B**_1_, …, **B**_L_]), where **β** (R x 1) is the (coefficient) vector of association between the response and the scores, and is estimated using least squares i.e. **β** = (**A**^*T*^**A**)^−1^**A**^*T*^**y**.

We therefore introduce a similar sum of squares term 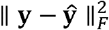 where **ŷ** = **Aβ**. This fit to the response **y** is restricted through a joint penalty on the coefficients factor *β*_*r*_ and the corresponding loadings **b**_:,*r*_ via the product 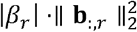. Here, **b**_:,*r*_ collects the *r*-th loadings across all features (concatenating across *L* blocks; i.e., ignoring the CLD structure), By multiplying | β_*r*_| and 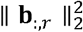 the penalty links each regression coefficient to the magnitude of its associated factor loadings, therefore, if a factor is important for **y** (i.e., *β*_*r*_ is large) the penalty also prevents unbounded growth in 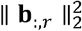. If *β*_*r*_ is zero, the loadings are not penalised by the **y**-side, allowing that factor to remain purely for describing **X**, making the full PESCAR loss:

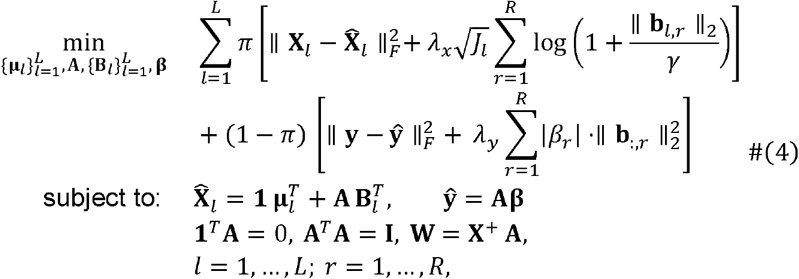

There is now a drive to fit a response **y** through a PCovR-like model (22). The balance between fitting **X** and **y** is controlled by the parameter *π*. In equation (4), **W**is introduced for cross-validation and prediction, and is estimated at each iteration *k* to enable **A** calculation from new or partially observed samples. Note that loss functions in equations (2) and (4) have been simplified, however in both cases they remain parametrised similar to (6) such that Bernoulli and Poisson distributed data blocks can be directly analysed (see supplementary section 4).

##### Feature Selection

We take the final scores **A** and loadings **B** and calculate a model-based association with **y**:

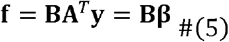

This provides a weighted average vector of loadings scaled by the coefficients (**β**), resulting in a scalar value for each feature that indicates similarity to **y** within the feature space of the model.

##### y-prediction

A new sample **x**_new_ is projected onto the score space via **w**, after which the resulting scores are multiplied by ***β*** to obtain *y*_new_

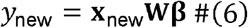

### Algorithm

The PESCAR algorithm takes as input a set of multi-block data matrices **X**_*l*_, a response vector **y**, regularisation parameters *λ*_*x*_, *λ*_*y*_, the GDP penalty parameter *γ*, block-specific_x_ scaling factors *α*_*l*_ (calculated using PCA as described in (6)), convergence criteria including maximum number of iterations and a tolerance for the loss function and *π* which controls the balance between exploratory and supervised analyses.

With these parameters set, initialisation involves generating starting values for the factor matrices **A**^0^ and 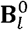 (via SVD, PLS or randomly), and the response coefficients ***β***^2^. The reconstructed block matrices 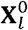 are formed and the initial loss is computed.

At each iteration, **A**^*k*^ is updated to **A**^*k*+1^ via a rank R SVD of the current reconstructed data 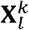 (i.e. the estimate of all L blocks concatenated multiplied by the current **B**^*k*^. Each **B**_*l*_ is then updated component-wise using a soft-thresholding operator that incorporates both the block-specific penalty and the response-related penalty, to form **B**_*l*_^*k*+l^. The reconstructed Matrices 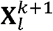 are then formed and the loss function is recomputed; convergence is assessed by the relative change in the loss. This process continues iteratively until convergence criteria are met or the maximum iteration number is reached. The algorithm returns the estimated factor matrices **A** and **B** reconstructed data matrices 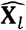, regression coefficients 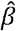, and the final loss value, outlined in Supplementary (section 2 Algorithm (1)).

### Model selection

In real data, because the underlying structure is unknown, we select the tuning parameters *λ*_*x*_ and *λ*_*y*_ by grid search, with a fixed value for *π*. For each fitted model we record the final reconstruction errors

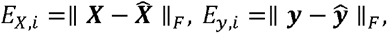

and retain only the Pareto-optimal models, i.e. those that are non-dominated: a model *h* is retained if there is no other model *g* such that *E*_*X,g*_ ≤*E*_*X,h*_ and *E*_*y,g*_ ≤ *E*_*y,h*_ with at least one strict inequality.

We normalise both errors to a common [0,1] scale and select the point closest to the Utopia point (the coordinate-wise minima of the two errors) using Euclidean distance. For an illustration of this procedure see figure 1 from (23).

Let *h* ∈ **ℋ** index the models on the Pareto front, and define the min–max scaled error coordinates

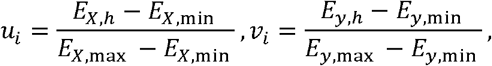

where the minima and maxima are taken over *i* ∈ **ℋ**. We then select

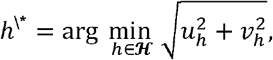

i.e. the Pareto-front model closest (in Euclidean distance) to the bottom-left corner (0,0) of the scaled error rectangle. This procedure is also used for simulation evaluation.

### Simulations

The following simulations characterise the operating range of PESCAR. Specifically, we assess robustness to the strength of ***y***-related structure within the ***X*** blocks, and to model rank mis-specification.

We create a rank 6 system across 3 **X** blocks with 2 Common, 1 Local (**X**_1_ and **X**_2_), and 3 Distinct components with **y** associated to the (weak) Local component i.e.

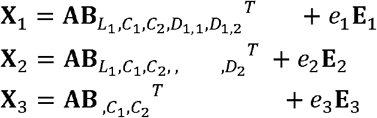

These components are weighted by **∑**_*X*_ ∈ ℝ^*L*×*R*^

Where:

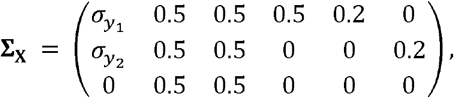

such that

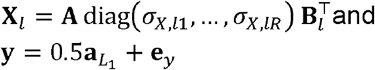

Where the six columns of **∑**_*X*_ correspond to components (*L*_1_, *C*_1_, *C*_2_,*D*_1,1_, *D*_1,2_, *D*_2_), and the three rows correspond to the blocks ***X***_**1**_,***X***_2_, and ***X***_3_. Nonzero entries indicate component presence within a block, and their magnitude determines signal strength. Furthermore, **A**^*T*^**A** = **I, 1**^*T*^**A** = 0, **B**, is sparse across columns i.e., features were simulated to load primarily on a single component, with limited overlap between components (*L*_1_, *C*_1_), (*C*_1_, *C*_2_), and (*C*_2_, *D*_1,1_). Noise matrices **E**_1:3_ are generated separately from random gaussian distributions, scaled such that they have a Frobenius norm of 1 and then rescaled by the scalars *e*_1:3_.

### Experiment 1: Noise

To investigate to what extent a signal, related to the supervision **y**, can be recovered from a noisy background we used the above backbone with 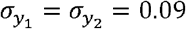, so that the combined size of the **y**-related signal was the smallest structured component in the data. Across 5 simulated datasets with random structure (i.e. randomised orthogonal **A** matrices), we systematically varied the noise level in all three data blocks jointly, setting *e*_1_ = *e*_2_ = *e*_3_ ∈ {0.6,0.9,1.2}, equating to designed noise levels of 30.61%, 39.82% and 46.88% in the simulated dataset.

### Experiment 2: Model misspecification

Since the underlying dimensionality of a real biological system is never known *a priori*, we investigated how model misspecification can affect the analysis.

#### Experiment 2a: Over-specification

To study over-specification, we used the same simulation framework as in Experiment 1, but fitted models with 7 components, thereby deliberately exceeding the true underlying rank. To assess whether PESCAR could still recover the weak *y*-related structure under over-specification, we systematically varied the size of the *y* -related component embedding by setting 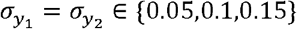. Noise levels were fixed at a realistic level of 0.15 (or 9.93% of the designed variation).

#### Experiment 2b: Under-specification

To assess the effect of rank under-specification, we examined whether supervision could enable recovery of a weaker ***y***-related component in the presence of stronger components, in particular by favouring it over one of the more strongly embedded **X** components. We repeated the setup from Experiment 2a, including the same *y*-embedding levels 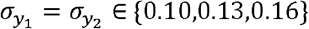 and fixed noise level of 0.15, but fitted models with 5 components rather than 7, thereby underestimating the true rank.

### Model Interpretation Simulation

In order to clearly demonstrate the interpretation of model outputs we simulated a new we simulated a rank-6 system across three predictor blocks with two Common components (*C*_1_, *C*_2_), two Local components (*L*_1_, *L*_2_), and two Distinct components (*D*_1_, *D*_2_), where *L*_1_ is shared by **X**_1_ and **X**_3_, *L*_2_ is shared by **X**_2_ and **X**_3_, *D*_1_ is specific to **X**_l_ and *D*_2_ is specific to **X**_2_.

Here all component block contributions are equal:

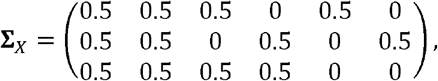

The six columns of **∑**_*X*_ correspond to (*C*_1_, *C*_2_,*L*_1_, *L*_2_,*D*_1_, *D*_2_), and the three rows correspond again to (**X**_1_, **X**_2_, **X**_3_). The block membership and overlap structure of the loading vectors were defined by a fixed component graph.

Within each block, features were simulated to load either on a single active component or on a pair of active components connected by an edge in the graph. Here, the edge topology was defined as:

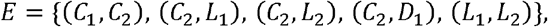

Hence, the graph determined the sparse support pattern and overlap structure of the loading matrices, and **∑**_*X*_ determined the magnitude of the corresponding rank-one signals. For features assigned to an edge (*r, s*), the paired nonzero loadings (*b*_*l,jr*_, *b*_*l,js*_)were generated according to an edge label, with labels corresponding to qualitatively different two-component relationships, these two dimensional patterns were: same values, spiral, negatively correlated, spiral and positively correlated (respectively to the pairs in *E*). Noise matrices **E**_l_, **E**_2_, and **E**_3_ were generated similarly to the above with e1 = e2 = e3 = 0.2.

The response in this case was generated to connect to two components as:

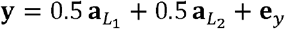

The simulated loading matrix here is represented as a hypergraph in **Fig. 1**.

### Performance evaluation

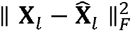

To evaluate performance, we calculate:

1. how much the estimated scores differ from the true scores i.e., the size of the residual of the estimated scores 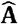 after orthogonal projection onto the column space of the true scores **A**_*true*_:

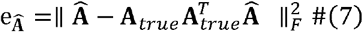
2. how good the estimate of the **y** related component is i.e., the size of the residual of the true **y** related component after orthogonal projection onto the column space of the estimated scores:

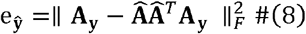
3. how many **y**-related features are properly recovered i.e., the geometric mean rank of *J*_target_ after feature selection:

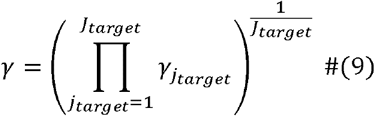

Lower values for all 3 metrics indicate better performance. The lower bound for 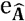 and e_ŷ_ is 0, the lower bound for *γ* is dependent on the number of **y** related features *J*_target_

### Real data

The example biological dataset is derived from a multi-omics study (20) in which the root transcriptome, metabolome and microbiome of tomato were analysed in a full factorial design with two growth conditions (N+/N-) and three substrates soil, gamma-irradiated (sterile) soil and rockwool) across four time points (4, 8, 12 and 16 days post treatment). At each combination of growth condition, substrate and time point, roots of five plants and their root exudate were collected. The transcriptome, metabolome and microbiome data were generated and pre-processed to approximately normal distributions for each block in line with the preprocessing strategies in the original paper (20).

## Results

### Experiment 1: Noise

We investigated how PESCAR compares to its unsupervised counterpart PESCA in recovery of **y** related features which are embedded as a small local component in **X**. Within realistic ranges of noise, the performance of supervised and unsupervised approaches was comparable at total subspace capture (top row ***Fig 2***.). Supervision helped to reduce the error in finding the **y** related component (middle row) if the noise in the data becomes larger and this also translated to better feature selection performance (bottom row).

**Figure 2.**
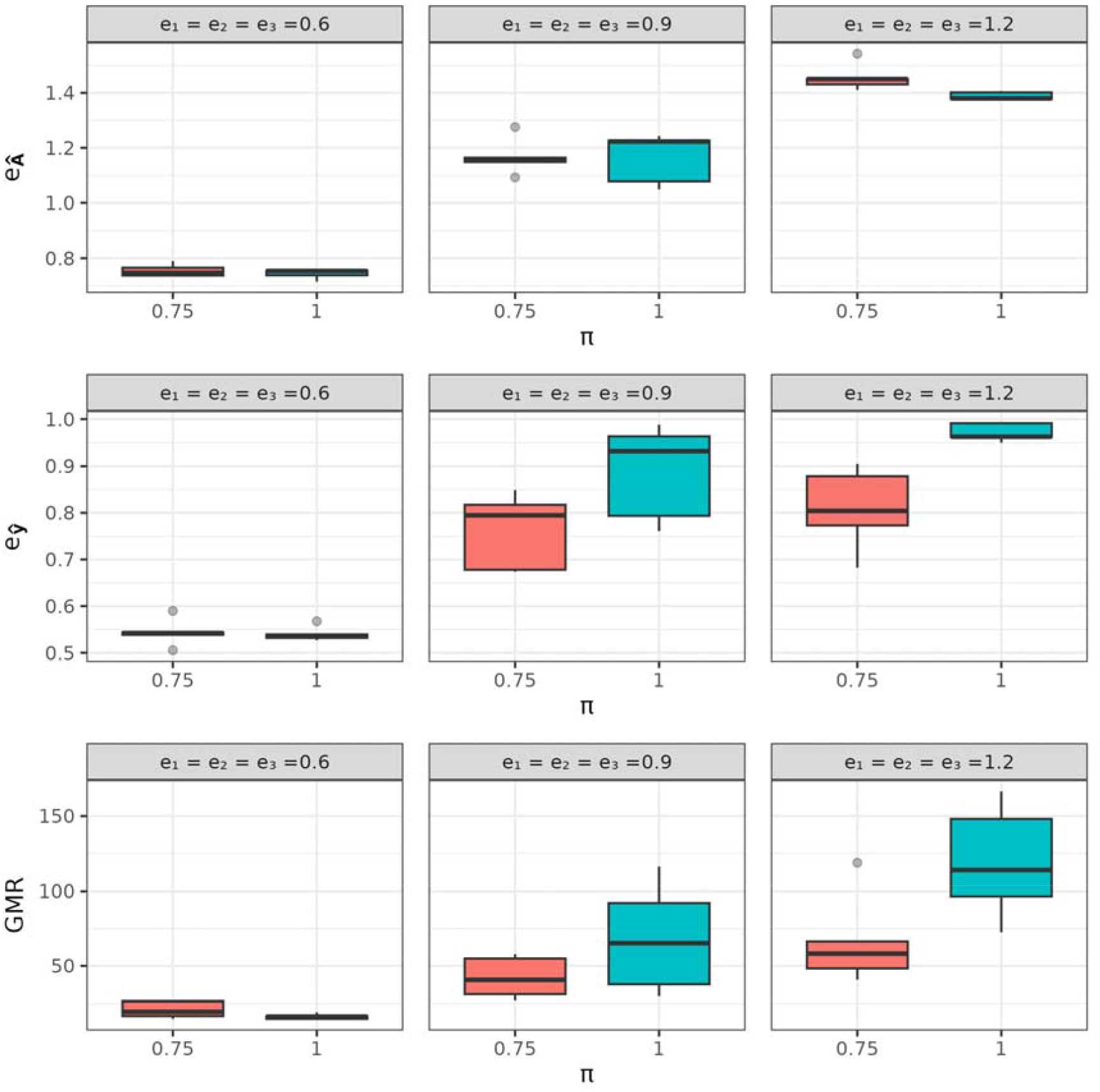
The effect of noise. Top row: Frobenius norm of 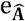, Middle row: Frobenius norm of e_ŷ;_. Bottom row: geometric mean rank *γ*. X axes are split by π values; 1 indicates no supervision i.e., original PESCA. Panel titles e1, e2 and e3 indicate per block noise levels.

**Figure 3.**
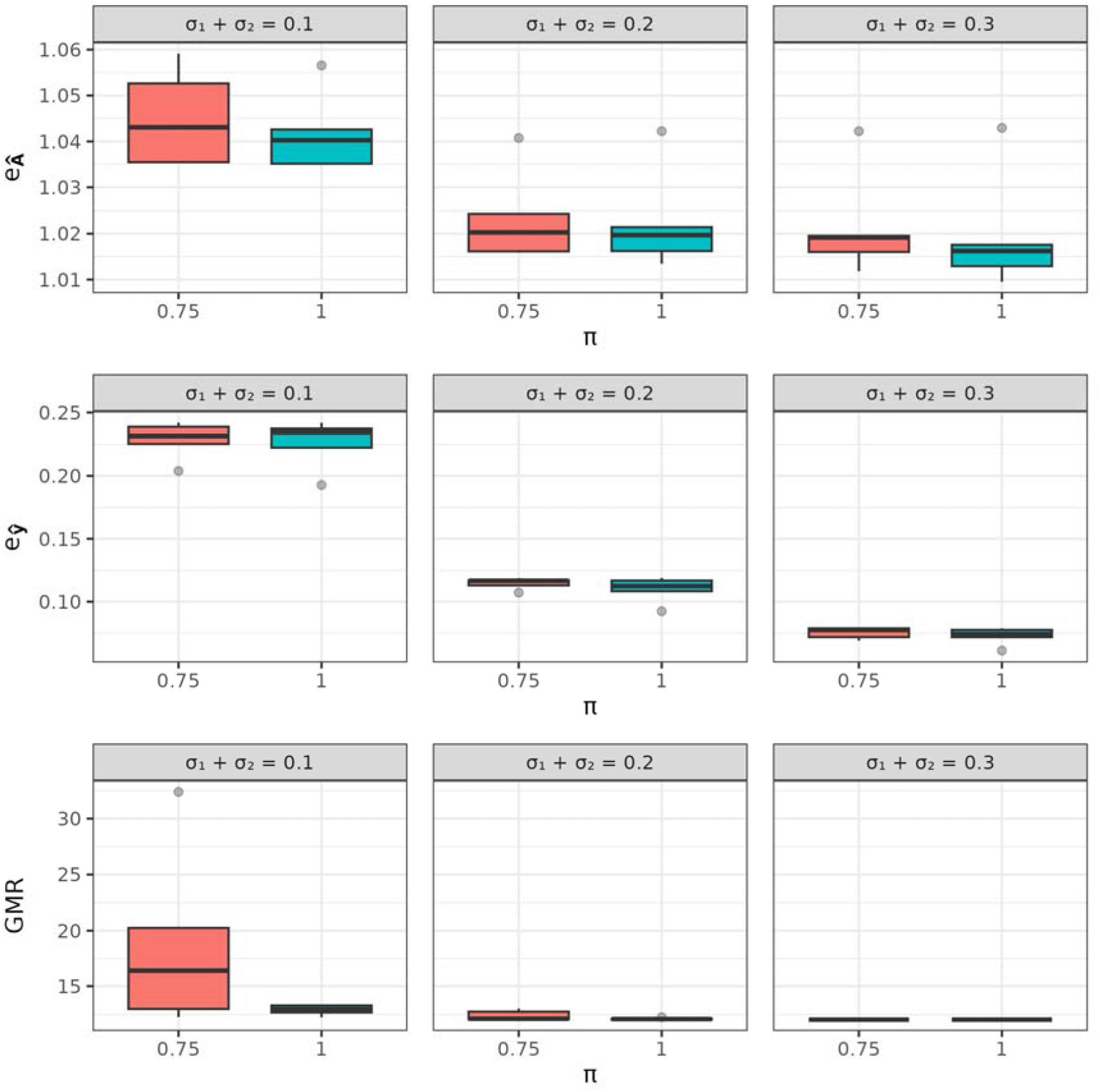
Effects of over-specification. Top row: Frobenius norm of 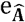, Middle row: Frobenius norm of e_ŷ_. Bottom row: geometric mean rank *γ*. X axes are split by π values; 1indicates no supervision i.e., original PESCA. Panel titles indicate strength of the embedding of the **y** related component as sum of Frobenius norms.

Since the rank of the system estimate is rarely known *a priori*, it is important to investigate how model misspecification affects integration results.

### Experiment 2a: Over specification

To investigate over-specification, a similar experiment to the above was carried out, however here the strength of the embedding of the **y** related component was varied and models of rank 7 were calculated. We saw mostly similar results to experiment 1: a slight breakdown at very high noise levels in both methods, but otherwise largely comparable results.

### Experiment 2b: Under specification

From a biological standpoint, multi-omics systems are shaped by many partially independent processes (e.g., pathway modules, environmental responses, developmental programs, and microbial guilds), each contributing their own sources of variation. In a multiblock setting, variation is further decomposed into CLD structure, which increases the effective dimensionality. Therefore, while the true rank is still expected to be far smaller than the number of measured features, it is not generally realistic to assume that a very small number of components captures the full biological system. However, the number of components that are directly important for a biological question are preferably small, since then considerably fewer candidates need to be physically investigated. To that end we postulate that model under-specification is the most likely case to occur in practice. To investigate the effect of under-specification, we conducted a similar experiment as experiment 2, and calculated models of rank 5. Results of this experiment showed that the supervised version had a slightly larger error in total subspace reconstruction (top row ***Fig 4***.). This is expected, as the aim is to use supervision to find a smaller component that describes **y**, whilst subject to model under specification this must happen at the expense of a larger component that describes only **X**. This component replacement postulation is clearly confirmed (middle row ***Fig 4***.) and propagates to feature selection improvements, too.

**Figure 4.**
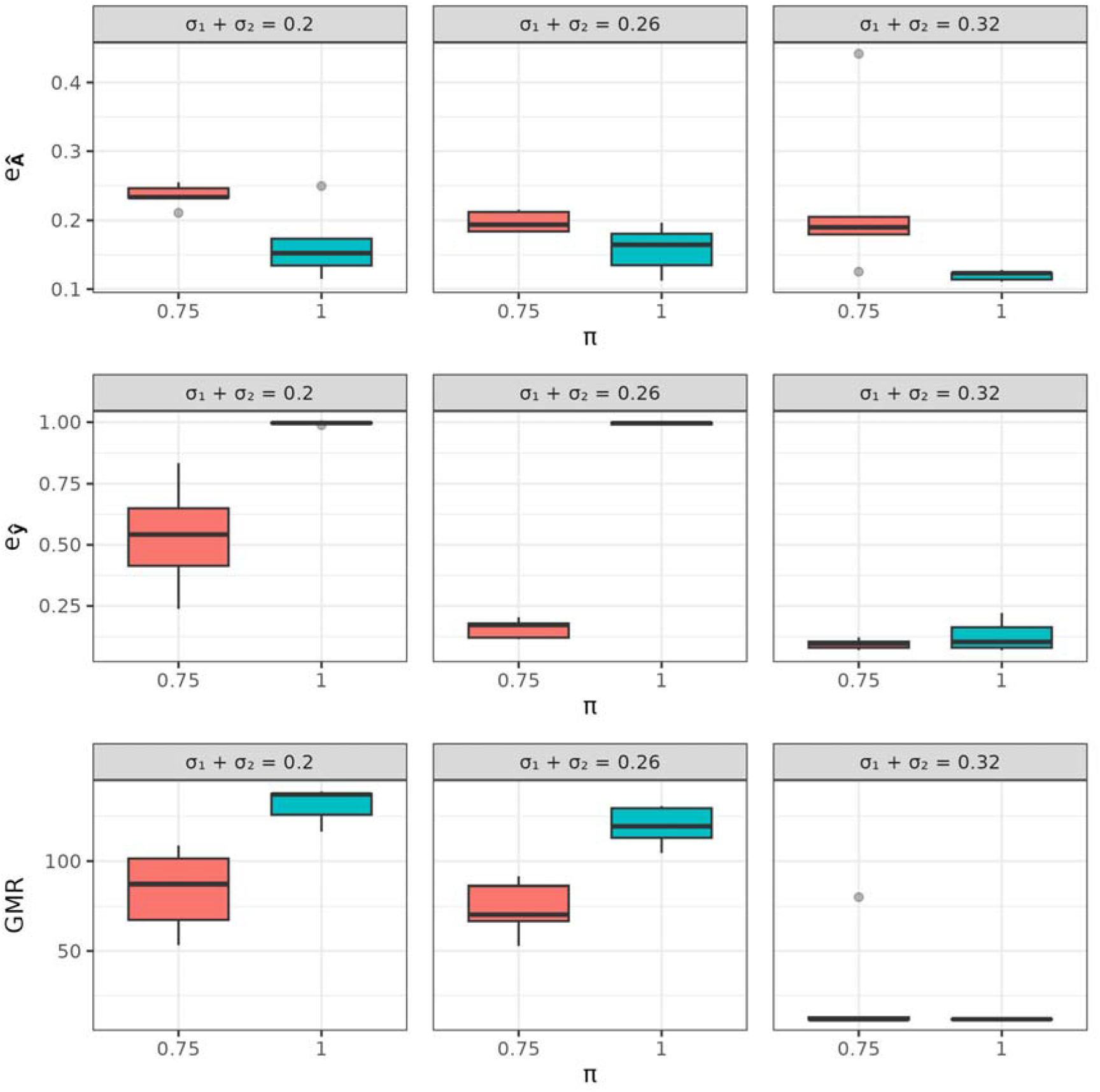
Effect of under-specification. Top row: Frobenius norm of 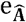, Middle row: Frobenius norm of e_ŷ_. Bottom row: geometric mean rank *γ*. X axes are split by π values; 1 indicates no supervision i.e., original PESCA. Panel titles indicate strength of the embedding of the **y** related component as sum of Frobenius norms.

### Model Interpretation: Simulated example

In the above simulations we have shown the use case for PESCAR. Here we demonstrate the model interpretation on an example simulated dataset. The primary outputs describe the relative contribution of each block to each component and indicate the loading profiles of the top features from each block as selected by equation (5). The model selected through Pareto front on the grid search of *λ*_*x*_ and *λ*_*y*_ (***Fig. 5A***) identifies 2 common components C1 and C2, 1 local component between blocks X1 and X3; C3, 1 local component between blocks X2 and X3; C4 and 2 distinct components (1 for block X1; C6 and 1 for block X2; C5, seen in the radar plot and full model loading profiles ***Fig. 5B-C***).

**Figure 5.**
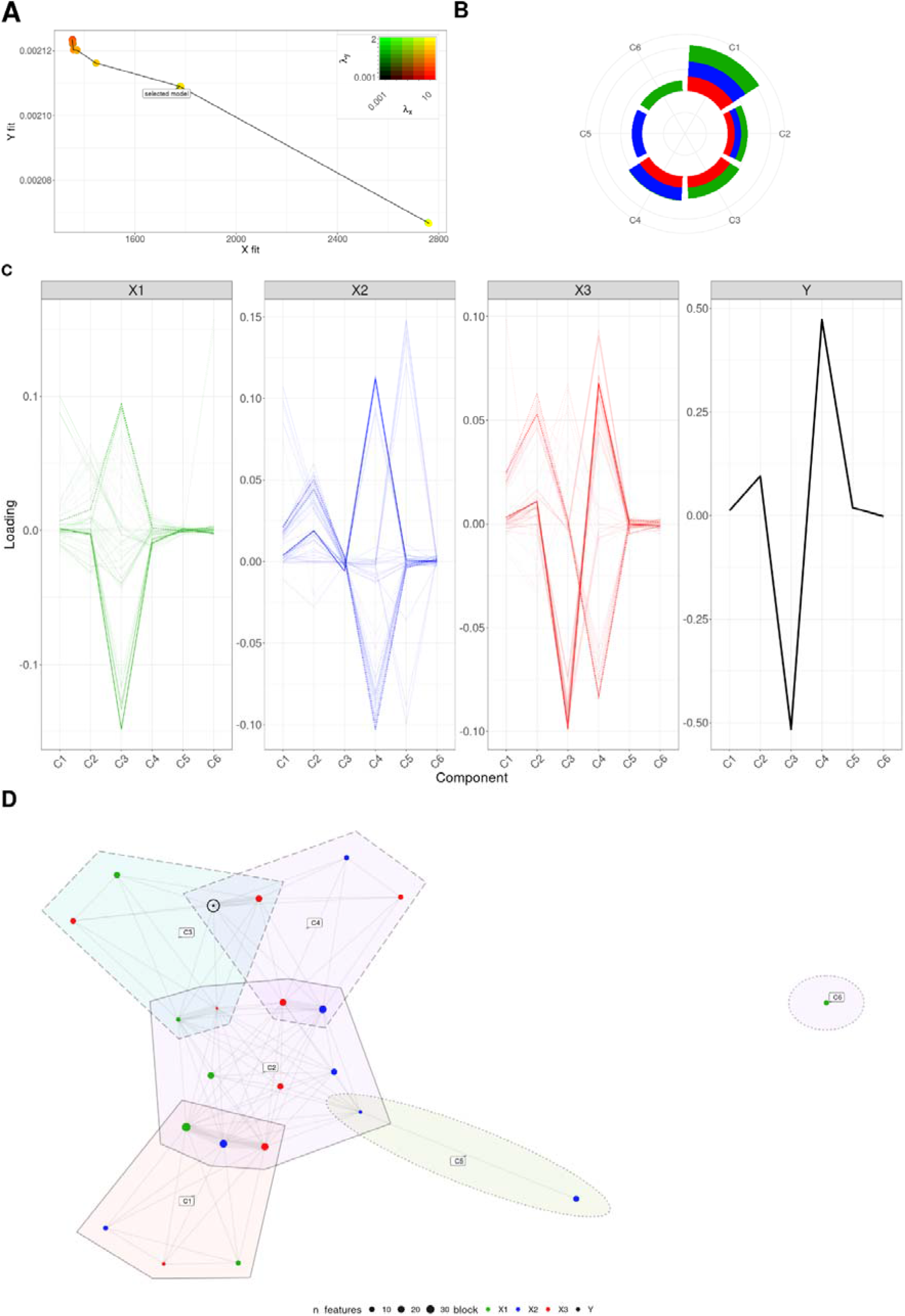
Model interpretation on example simulated dataset. **A**; Model selection based on the Pareto front over the *λ*_*x*_ and *λ*_*y*_ grid search, indicated in plot inset; redness corresponds to *λ*_*x*_, greenness corresponds to *λ*_*y*_, yellow indicates both penalties in use. **B**; Radar plot showing the variance explained by each component in each block, used to identify common, local, and distinct component structure. **C**; Block-wise loading profiles of the selected model across all components, showing the feature loading patterns within each block, line opacity scaled by **y**-relatedness (f). **D**; Hypergraph representation of full model derived from overlap in the nonzero loading support across components, summarising the higher-level topology of the fitted model, edge width indicates the number of components that a pair of nodes are both nonzero.

From the loading profile line plots it is evident that the supervision vector (“Y”) is primarily related to C3 and C4 with an overall loading signal profile that relates most strongly to a set of features in X3 (on both C3 and C4) and also partially to sets of X1 and X3 features in C3 and sets of features from X2 and X3 in C4. These sets of features display clearly different behaviours across the model as a whole, and thus can be interpreted as being linked to Y through different parts of the process being measured. In the loading profile line graphs, we can also see other clusters of features that display alternative profiles. These profiles are not directly connected to the response (no/limited signal on C3 or C4) but interesting to note as contributing to the overall topology of the dataset, through their modelled covariance structure. For example the sets of features with high negative values on C4 in X2 and X3 also have high positive values on C2. This type of interpretation can be translated as Y related and adjacent pathways. Except for C6, which is distinct to block X1, the recovered components form a single connected graph through indirect overlap. This higher-level topology, defined by overlap in the loading matrix, is shown in ***Fig. 5D***. The reconstructed hypergraph has the same topology as the input (***Fig. 1***), the one difference is the node sizes, this indicates that a small number of features were not recovered at some nodes, due to the added noise added and penalisation.

### Real Data

To demonstrate the performance of PESCAR on a real multi-omics data set we used experimental data from (20). The experiment conducted involved tomato plants which were subjected (or not) to nitrogen deficiency and grown on three different substrates. Transcriptome, metabolome (root exudate and root extract, negative and positive LCMS mode) and 16S root microbiome data were generated on this material, creating 6 blocks. In the experiment it was demonstrated that N deficiency resulted in the production of a signalling molecule (strigolactone) which was used here as **y** to find features across the omics datasets that may contribute to its formation or respond to its level (primary **y**-related candidates), while taking into account that other processes contribute to other components.

For this dataset, we first fitted a 20-component PESCAR model and subsequently refitted the model with 10 components, 10 components in the original model each explained less than 1% of the total variation. The final 10-component model, selected using Pareto front model selection (**Fig. 6A**), comprised 5 common components and 5 local components; the local components did not involve the exudate metabolome data blocks (**Fig. 6B-C**). The response variable, corresponding to the strigolactone level, was most strongly associated with C1 and C2. Based on this response profile, features most strongly associated with the response were quantified, and top candidate biological processes underlying this response-associated variation were identified using the feature selection procedure described in equation (5). These top candidates are discussed at a functional level below. By contrast, the hypergraph visualisation (**Fig. 6D**) was constructed from a component-wise thresholding of the full loading matrix and therefore reflects broader component support structure across the model, in order to contextualise the **y**-related signal in the larger system. As such, the figure includes a subset of the primary -related candidates, but is not restricted to them. For each component, the 25 non-zero **X**-feature loadings with the largest absolute values were retained. The resulting hypergraph suggests two broad groups of features: one composed almost entirely of metabolic features, with no genes and only a single ASV, and another with a more balanced mixture of features from all blocks. This pattern is consistent with a distinction between on the one hand more internal regulation and on the other a biosynthetic state of the tomato root and active chemical shaping of the rhizosphere. Top **y**-related candidate functions are briefly discussed below (full details with annotations in dataset **S1**).

**Figure 6.**
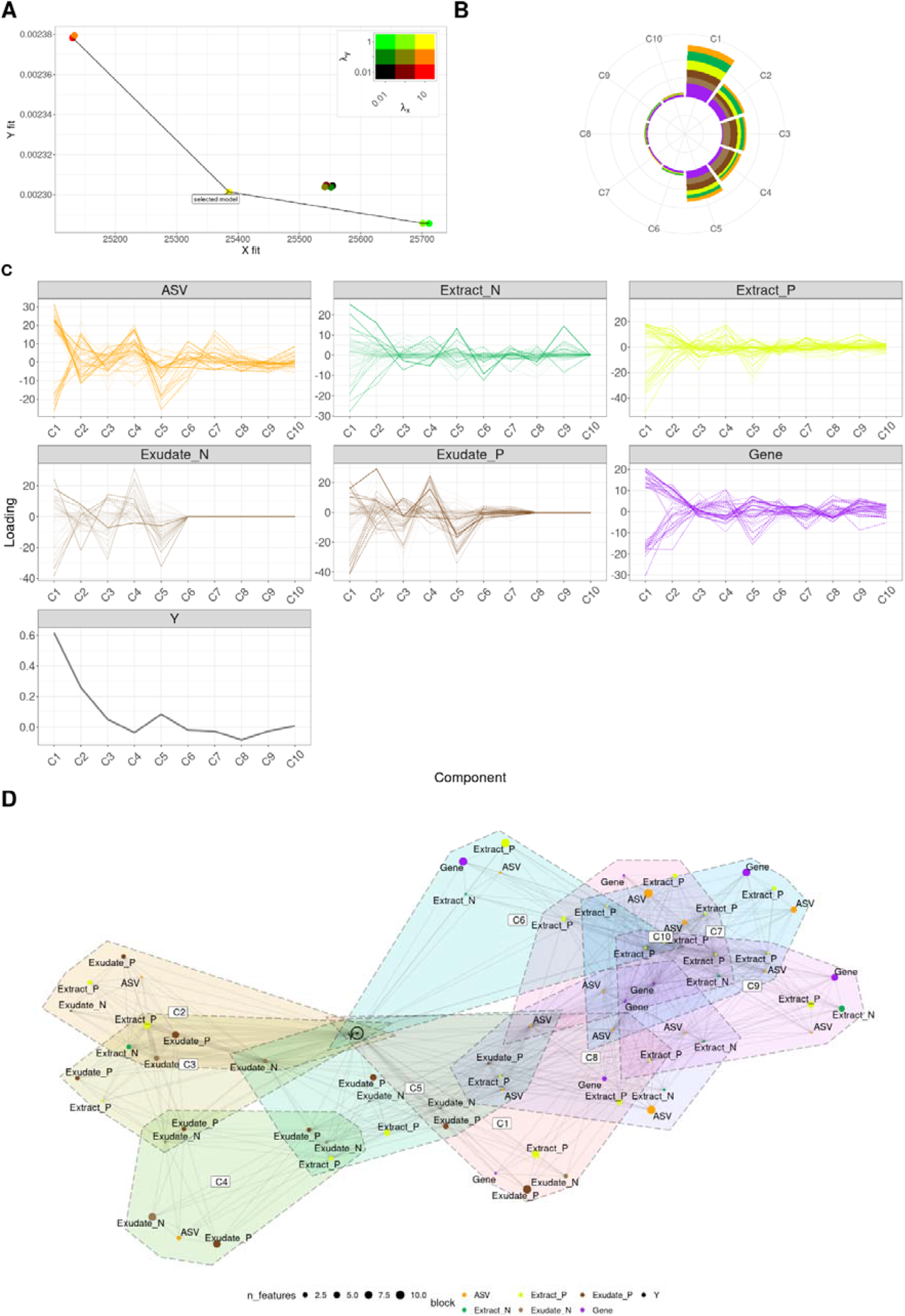
demonstration of PESCAR on real data. **A**; Model selection based on the Pareto front over the *λ*_*x*_ and *λ*_*y*_ grid search, indicated in plot inset; redness corresponds to *λ*_*x*_ greenness corresponds to *λ*_*y*_, yellow indicates both penalties in use. **B**; Radar plot showing the variance explained by each component in each block, used to identify common, local, and distinct component structure. **C**; Block-wise loading profiles of the selected model across all components, showing the feature loading patterns within each block, line opacity scaled by **y**-relatedness (f). **D**; Hypergraph representation of the full model derived from overlap in the nonzero loading support across components, summarising the higher-level topology of the fitted model, node size indicates number of features represented, edge width indicates the number of components that a pair of nodes are both nonzero.

*Genes;* top candidate genes indicate a combined signal of defence, lipid/carotenoid-derived signalling, nutrient homeostasis, redox regulation, and kinase-mediated regulation. In particular Cytochrome p450s (`Solyc10g018150.2.1`, `Solyc08g079300.3.1`) as well as a known strigolactone precursor (`Solyc01g005940.3.1`) were detected as strongly related to the strigolactone response variable, and several defence related genes were detected as having a strong opposing signature including pathogenesis related (`Solyc00g174340.2.1`), nitrate transporter (`Solyc08g007430.2.1`) and MAPKK (`Solyc01g111880.3.1) among others.`

*Metabolites – Extract;* the extract-associated metabolites comprised alkaloids, amino acids and peptides, polyketides, fatty acids, carbohydrates, butenolides, glycosyl compounds, spirosolanes and derivatives, and alkyl phosphates.

*Metabolites – Exudate;* the exudate-associated metabolites comprised shikimates and phenylpropanoids, alkaloids, carbohydrates, terpenoids, fatty acids, polyketides, and amino acids and peptides, indicative of rhizosphere-facing nutrient, signalling, and defence chemistry.

*Microbes;* the top candidate taxa were dominated by plant-associated, nitrogen-cycling (Rhizobium), plant growth promoting (Pelomonas) and known plant symbionts/strigolactone mediated helpers (Devosia).

Together, these functions indicate that the recovered axis reflects coordinated root-internal regulation, exudate chemistry, and rhizosphere microbial structure, within which strigolactone-related biology appears to play a prominent role. These results are mostly consistent with the results from (20).

## Discussion and conclusions

In this work, we introduced PESCAR, an extension of PESCA (6) designed to uncover common local and distinct (CLD) structures in multi-block data using the information in a response variable to (partially) shape the model. One of the key advantages of PESCAR is its ability to identify smaller sources of information in the data of specific interest that are missed using unsupervised PESCA. These subtle, response-related components can be critical for understanding the underlying processes, particularly when the signal of interest is weak or hidden by noise or other more variant components.

Our simulation studies confirm the effectiveness of PESCAR in recovering these smaller, response-related components. Across a range of parameters, PESCAR consistently identified relevant CLD structures. Application to a real-world dataset (20) further supports the utility of PESCAR. In a multi-omics dataset, PESCAR uncovered biologically meaningful patterns aligned with documented functional relationships, notably recovering known elements of the strigolactone pathway related to the response variable used for supervision.

In many latent variable models, components are treated as independent objects and interpreted one at a time (e.g., “genes 1–5 define component 1, representing process A”). Orthogonality of the score matrix is mathematically convenient and simplifies estimation ((24),(25)). However, orthogonality in the latent space does not imply that the underlying biology is independent or separable into non-interacting processes. In reality, biological mechanisms are coupled, and their molecular signatures can and do overlap across pathways, blocks, and measurement layers. In our framework, we therefore keep orthogonality in the scores, but allow and investigate overlap in feature support across components through the loading matrix. Concretely, the same feature can contribute to multiple components, which enables a more system-level interpretation: we can study not only “which features load on which component”, but also how sets of features co-occur across components and blocks. This is exactly the structure exploited by our topology-oriented summaries (hypergraph visualisations), where edges/hyperedges are defined by shared support patterns. This provides a more system-level interpretation, however the orthogonality assumption itself remains a simplifying constraint.

Allowing overlap in the loadings can exacerbate rotational non-uniqueness in the loading space, because multiple rotated bases may provide nearly equivalent fit while redistributing feature support across components. In other words, the same underlying structure can be viewed from different orientations, each giving a similar overall picture but assigning features somewhat differently across components. This is further exacerbated when the sizes of the components are similar (see supplementary section 3 for a mathematical description on why this is not an issue for the purposes of sample-sample relationships and the corresponding covariance of features, as well as for prediction). In our experiments, this did not hinder the primary goal of identifying response-associated structure and cross-block feature associations. However, it can complicate exact recovery of the simulated ground-truth CLD topology when CLD is defined as a binary component-block membership pattern. This suggests that, under realistic overlap, “topology recovery” is better posed as recovering degrees of shared signal rather than exact binary memberships.

There are still some limitations to the current approach. Component subscription overlap can be represented in the loadings, but the orthogonality assumption itself remains a limitation of the current framework. While orthogonal components simplify the mathematical framework, biological processes as investigated in (20) are rarely strictly independent. Imposing orthogonality may therefore obscure or distort some relationships between processes. Future work could explore relaxing this constraint, by introducing a soft orthogonality penalty. More generally, orthogonality may also limit downstream analyses aimed at studying directed dependence between components, such as component-on-component regression, although the mixed component membership model we introduce here introduces intermediate feature sets that can circumvent this.

Future work in this line should consider new definitions of CLD structure, perhaps not as a binary matrix, indicating a block’s subscription to a component, but rather a quantifiable degree of signal overlap across components and blocks, and topology summaries that are robust to (admissible) rotations. Note, the CLD topology recovered here by PESCAR should not be interpreted as a causal graph. Rather, a summary of structured overlap in (response-relevant) signal across components and blocks. In this sense, the hypergraph representation is best viewed as a hypothesis-generating tool: it highlights candidate groups of features and modalities that vary together in (a part of) the model, but it does not by itself determine directionality or mechanism. Establishing causal relationships would require additional assumptions or sources of information, such as temporal ordering, perturbation experiments, or prior biological knowledge.

Overall the findings in this work suggest that integrating response information into the estimation of CLD structure can improve interpretability and facilitate hypothesis driven exploration in complex biological systems.

In summary, PESCAR extends PESCA by jointly estimating CLD structure and exploiting response information during fitting. This coupling helps reveal ***y***-aligned components that can be hidden in unsupervised decompositions and provides a structured basis for interpreting multivariate relationships across multi-block data in complex systems. Code and data for this project are available at https://github.com/BiosystemsDataAnalysis/PESCAR.

## Acknowledgements

We acknowledge funding by the Dutch Research Council (NWO/OCW) for the MiCRop Consortium program, Harnessing the second genome of plants (Grant number 024.004.014; to HB, LD, AKS, JAW), the Dutch Research Council (NWO-TTW grant 16873 Holland Innovative Potato; to HB,LD and FW), Research Priority Area on Personal Microbiome Health of the University of Amsterdam (to GRvdP) and the Data Science Centre of the University of Amsterdam (to FW).

