## supplementary for "Supervised restricted data fusion with common, local & distinct components"

**PESCAR – Supplementary Materials**

**1. Loss Function Gradient**

Since the concatenation: $\parallel\mathbf{b}_{:,r}\parallel_{2}^{2} = \sum_{l=1}^{L} \parallel\mathbf{b}_{l,r}\parallel_{2}^{2}$ and, in implementation, $\boldsymbol{\beta}$ is re-estimated by ordinary least squares on the current $\mathbf{A}^{k}$; the fit term for $\mathbf{y}$ does not affect the $\mathbf{B}$-update step except through the current $\boldsymbol{\beta}$ values. Therefore, holding $\mathbf{A}$ and $\boldsymbol{\beta}$ fixed during the $\mathbf{B}$-step, the $\mathbf{B}$-dependent supervised term reduces to $\left( 1-\pi\right) \lambda_{y}\sum_{r=1}^{R} \left| \beta_{r} \right|\sum_{l=1}^{L} \parallel\mathbf{b}_{l,r}\parallel_{2}^{2}$, note that we are able to use the block specific notation here which enables more succinct gradient derivation.

The blockwise gradient of the loss function with respect to $\mathbf{b}_{l,r}$ is

$$\begin{matrix} \nabla_{\mathbf{b}_{l,r}}\left( \left( 1-\pi\right)\lambda_{y}\left| \beta_{r} \right|\sum_{l} \parallel\mathbf{b}_{l,r}\parallel_{2}^{2} \right) = 2 \left( 1-\pi\right)\lambda_{y}\left| \beta_{r} \right| \mathbf{b}_{l,r} \end{matrix}$$

Define the component-wise ridge coefficient (shared across all blocks l for a fixed r):

$c_{r}^{\left( k \right)} = \left( 1-\pi\right) \lambda_{y} \left| \beta_{r}^{\left( k \right)} \right|$ and the group lasso threshold $\tau_{l,r}^{\left( k \right)} = \pi\lambda_{x} \omega_{l,r}^{\left( k \right)}$.

Where $\omega_{l,r}^{\left( k \right)} = \frac{1}{\gamma+\parallel\mathbf{b}_{l,r}^{\left( k \right)}\parallel_{2}}$ .

The Frobenius norm loss can be rewritten as:

$$\left\| {\hat{\mathbf{X}}}_{l}-\mathbf{A}\mathbf{B}_{l}^{\top} \right\|_{F}^{2}=tr\left( \left( {\hat{\mathbf{X}}}_{l}-\mathbf{A}\mathbf{B}_{l}^{\top} \right)^{\top}\left( {\hat{\mathbf{X}}}_{l}-\mathbf{A}\mathbf{B}_{l}^{\top} \right) \right),$$

so

$$\left\| {\hat{\mathbf{X}}}_{l}-\mathbf{A}\mathbf{B}_{l}^{\top} \right\|_{F}^{2}=tr\left( {\hat{\mathbf{X}}}_{l}^{\top}{\hat{\mathbf{X}}}_{l} \right)-2 \mathrm{tr}\left( \mathbf{B}_{l}\mathbf{A}^{\top}{\hat{\mathbf{X}}}_{l} \right)+tr\left( \mathbf{B}_{l}\mathbf{A}^{\top}\mathbf{A}\mathbf{B}_{l}^{\top} \right).$$

Since $\mathbf{A}^{\top}\mathbf{A}=\mathbf{I}$, this becomes

$$\left\| {\hat{\mathbf{X}}}_{l}-\mathbf{A}\mathbf{B}_{l}^{\top} \right\|_{F}^{2}=2 \mathrm{tr}\left( \mathbf{B}_{l}\mathbf{A}^{\top}{\hat{\mathbf{X}}}_{l} \right)+\left\| \mathbf{B}_{l} \right\|_{F}^{2}.$$

With the non B related terms dropped. Now define

$$\mathbf{Z}_{l}:={\hat{\mathbf{X}}}_{l}^{\top}\mathbf{A}.$$

Then

$$\mathrm{tr}\left( \mathbf{B}_{l}\mathbf{A}^{\top}{\hat{\mathbf{X}}}_{l} \right)=tr\left( \mathbf{B}_{l}\mathbf{Z}_{l}^{\top} \right),$$

and therefore

$$\left\| {\hat{\mathbf{X}}}_{l}-\mathbf{A}\mathbf{B}_{l}^{\top} \right\|_{F}^{2}=\left\| \mathbf{Z}_{l}- \mathbf{B}_{l} \right\|_{F}^{2} up to terms independent of \mathbf{B}_{l}$$

Let $\mathbf{z}_{l,r} := {\hat{\mathbf{X}}}_{l}^{\top} \mathbf{a}_{r}$. The MM surrogate (data fit and linearised concave penalty) plus the quadratic coupling yields, for each (l,r), the convex subproblem:

$$\begin{matrix} \min_{\mathbf{b}\in\mathbb{R}^{j_{l}}} \frac{1}{2} \parallel\mathbf{z}_{l,r}-\mathbf{b}_{l,r}\parallel_{2}^{2} + \tau_{l,r}^{\left( k \right)} \parallel\mathbf{b}_{l,r}\parallel_{2} + c_{r} \parallel\mathbf{b}_{l,r}\parallel_{2}^{2}. \end{matrix}$$

First, expanding the squared norm term:

$$\begin{matrix} \left\| \mathbf{z}_{l,r}-\mathbf{b}_{l,r} \right\|_{2}^{2} & =\left( \mathbf{z}_{l,r}-\mathbf{b}_{l,r} \right)^{\top}\left( \mathbf{z}_{l,r}-\mathbf{b}_{l,r} \right) \\ & =\mathbf{z}_{l,r}^{\top}\mathbf{z}_{l,r} - 2 \mathbf{z}_{l,r}^{\top}\mathbf{b}_{l,r} + {\mathbf{b}_{l,r}}^{\top}\mathbf{b}_{l,r}. \end{matrix}$$

The gradient of the differentiable/smooth part can be calculated as:

$$\begin{matrix} \nabla_{\mathbf{b}_{l,r}}f\left( \mathbf{b}_{l,r} \right) & =1/2 \nabla_{\mathbf{b}}\left( \mathbf{z}_{l,r}^{\top}\mathbf{z}_{l,r} \right) - 1/2 \nabla_{\mathbf{b}}\left( 2 \mathbf{z}_{l,r}^{\top}\mathbf{b}_{l,r} \right) + 1/2 \nabla_{\mathbf{b}}\left( {\mathbf{b}_{l,r}}^{\top}\mathbf{b}_{l,r} \right) + \nabla_{\mathbf{b}}\left( c_{r} {\mathbf{b}_{l,r}}^{\top}\mathbf{b}_{l,r} \right) \\ & =0 - 1/2 \left( 2 \mathbf{z}_{l,r} \right) + 1/2 \left( 2 \mathbf{b}_{l,r} \right) + 2c_{r} \mathbf{b}_{l,r} \\ & =-\mathbf{z}_{l,r} + \mathbf{b}_{l,r} + 2c_{r} \mathbf{b}_{l,r} \\ & =\left( 1+2c_{r} \right) \mathbf{b}_{l,r} - \mathbf{z}_{l,r}. \end{matrix}$$

$\nabla_{\mathbf{b}_{l,r}}\left[ 1/2\parallel\mathbf{z}_{l,r}-\mathbf{b}_{l,r}\parallel_{2}^{2}+c_{r}\parallel\mathbf{b}_{l,r}\parallel_{2}^{2} \right]=\left( 1+2c_{r} \right)\mathbf{b}_{l,r}-\mathbf{z}_{l,r}$.

Hence the first-order (subgradient) optimality condition is

$\mathbf{0} \in\left( 1+2c_{r} \right) \mathbf{b}_{l,r} - \mathbf{z}_{l,r} + \tau_{l,r}^{\left( k \right)} \partial\parallel\mathbf{b}_{l,r}\parallel_{2}$.

Solving the optimality condition gives a scaled group soft-threshold:

$$\begin{matrix} \mathbf{b}_{l,r}^{\left( k+1 \right)} = \left\{ \begin{matrix} \mathbf{0}, & \parallel\mathbf{z}_{l,r}\parallel_{2}\leq\tau_{l,r}^{\left( k \right)}, \\ \frac{1}{1+2c_{r}}\left( 1-\frac{\tau_{l,r}^{\left( k \right)}}{\parallel\mathbf{z}_{l,r}\parallel_{2}} \right) \mathbf{z}_{l,r}, & \text{otherwise.} \end{matrix} \right. \end{matrix}$$

This is exactly the original PESCA group soft-threshold with a component-wise scaling factor $1/\left( 1+2c_{r} \right)$.

For the $\mathbf{B}$ subproblem PESCA originally solves $\min_{\mathbf{b}} 1/2\parallel\mathbf{z}-\mathbf{b}\parallel_{2}^{2}+\tau\parallel\mathbf{b}\parallel_{2}$, with optimality condition $\mathbf{0}\in\mathbf{b}-\mathbf{z}+\tau\partial\parallel\mathbf{b}\parallel_{2}$ and solution $\mathbf{b}=\left( 1-\tau/\parallel\mathbf{z}\parallel_{2} \right)_{+} \mathbf{z}$. Our block concatenated supervised term adds the smooth gradient $2c_{r} \mathbf{b}$, modifying the condition to:
$\mathbf{0}\in\left( 1+2c_{r} \right) \mathbf{b}-\mathbf{z}+\tau\partial\parallel\mathbf{b}\parallel_{2}$, and hence the scaled solution above.

The GDP part is still handled by MM (linear upper bound via $\omega_{l,r}^{\left( k \right)}$), and the supervised coupling is convex and treated exactly. Therefore, each $\mathbf{B}$-update minimizes the per-block surrogate and the overall objective is non-increasing across $\mathbf{B}$-steps, this preserves monotone descent behaviour.

**2. PESCAR algorithm (supplementary algorithm 1)**

**Input:**

$$\{\mathbf{X}_{l}{\}}_{l=1}^{L}, \mathbf{y}, \{\alpha_{l}{\}}_{l=1}^{L}, \lambda_{x}, \lambda_{y}, \gamma, \text{maxit}, \text{tol\_obj}, \text{etc.}$$

**Output:**

$$\hat{\mathbf{A}}, \{{\hat{\mathbf{B}}}_{l}\}, \hat{\boldsymbol{\mu}}, \hat{\beta}$$

1. **Initialisation:**
   - Set iteration counter $k\leftarrow0$.
   - Initialise $\mathbf{A}^{0}\in\mathbb{R}^{I\times R}$ (using SVD, PLS or randomly).
   - Initialise each $\mathbf{B}_{l}^{0}\in\mathbb{R}^{J_{l}\times R}$ and $\boldsymbol{\mu}_{l}^{0}$.
   - Form $\mathbf{X}_{l}^{0}=\mathbf{1} \left( \boldsymbol{\mu}_{l}^{0} \right)^{T}+\mathbf{A}^{0} \left( \mathbf{B}_{l}^{0} \right)^{T}, l=1,\ldots,L.$
   - Calculate $\boldsymbol{\beta}^{\boldsymbol{0}}$ from $\mathbf{A}^{0}$.
   - Compute the initial loss $\mathcal{l}^{0}$.
2. **While not converged:**
   - For $k=0,\ldots,\text{maxit}-1$ do:
     - **(a) Update** $\boldsymbol{\beta}$
     - **(b) Update** $\mathbf{A}$**:**
       - Compute the rank $R$ SVD:

$${\hat{\mathbf{X}}}^{k} {\hat{\mathbf{B}}}^{k}=\mathbf{U} \mathbf{D} \mathbf{V}^{T}$$

- Set $\mathbf{A}^{k+1}\leftarrow\mathbf{U} \mathbf{V}^{T}$
- **(c) Update each** $\mathbf{B}_{l}$**:**
  - For $l=1,\ldots,L$ and each component $r=1,\ldots,R$:
    - $\sigma_{l,r}^{k}\leftarrow\parallel\mathbf{b}_{l,r}^{k}\parallel_{2}.$
    - $\omega_{l,r}^{k}\in\partial g\left( \sigma_{l,r}^{k} \right).$
    - $\tilde{\lambda}_{l,r}\leftarrow\lambda_{x} \sqrt{J_{l}} \omega_{l,r}^{k} {\alpha_{l}}/{\rho_{l}}.$
    - $\begin{matrix} \mathbf{b}_{l,r}^{\left( k+1 \right)} = \left\{ \begin{matrix} \mathbf{0}, & \parallel\mathbf{z}_{l,r}\parallel_{2}\leq\tau_{l,r}^{\left( k \right)}, \\ \frac{1}{1+2c_{r}}\left( 1-\frac{\tau_{l,r}^{\left( k \right)}}{\parallel\mathbf{z}_{l,r}\parallel_{2}} \right) \mathbf{z}_{l,r}, & \text{otherwise.} \end{matrix} \right. \end{matrix}$
  - Collect the columns $\mathbf{b}_{l,r}^{k+1}$ into $\mathbf{B}_{l}^{k+1}.$
- **(d) Form new** ${\hat{\mathbf{X}}}^{k+1}$ **and check convergence:**
  - ${\hat{\mathbf{X}}}^{k+1}\leftarrow\mathbf{A}^{k+1} \left( \mathbf{B}^{k+1} \right)^{T}$
  - Compute $\mathcal{l}^{k+1}$ using loss function.
  - If $\frac{\mathcal{l}^{k}-\mathcal{l}^{k+1}}{\left| \mathcal{l}^{k} \right|}<\text{tol\_obj}$, stop; otherwise, set $k\leftarrow k+1.$

3. **If converged:**

Return $\hat{\mathbf{A}}, \hat{\mathbf{B}}, \hat{\boldsymbol{\mu}}, \hat{\boldsymbol{\Sigma}}, \hat{\beta}, \mathcal{l,\ldots}$

**3. Rotational Freedom and uniqueness**

The estimated loading structure has two conceptually different forms of sparsity. First, there is sparsity, which induces the common, local, and distinct (CLD) structure by determining which blocks contribute to each component. Second, there is sparsity, that determines which features within a contributing block are active on that component. The first governs the broad multiblock topology of the solution, whereas the second governs the finer feature-level interpretation.

Consider any square orthogonal matrix $\mathbf{Q}\in\mathbb{R}^{R\times R}$. If the CLD block-support pattern is sufficiently restrictive to identify each component uniquely (up to trivial relabelling and sign changes), then no orthogonal rotation can be applied without altering that CLD structure. By contrast, if two or more components have compatible block-support patterns, so that the CLD constraints do not fully determine the basis, then an orthogonal re-basing within the corresponding latent subspace is possible.

Under such a re-basing,

$$\left( \mathbf{A},\mathbf{B} \right)\mapsto\left( \mathbf{A}_{\mathrm{rot}},\mathbf{B}_{\mathrm{rot}} \right):=\left( \mathbf{AQ}, \mathbf{BQ} \right).$$

Then

$$\mathbf{A}_{\mathrm{rot}}\mathbf{B}_{\mathrm{rot}}^{\top}=\left( \mathbf{AQ} \right)\left( \mathbf{BQ} \right)^{\top}=\mathbf{A}\left( \mathbf{Q}\mathbf{Q}^{\top} \right)\mathbf{B}^{\top}=\mathbf{A}\mathbf{B}^{\top}.$$

Hence the fitted reconstruction

$$\boldsymbol{\Theta}=\mathbf{1}\boldsymbol{\mu}^{\top}+\mathbf{A}\mathbf{B}^{\top}$$

and the associated data-fit term are unchanged.

The score Gram matrix is also invariant:

$$\mathbf{A}_{\mathrm{rot}}\mathbf{A}_{\mathrm{rot}}^{\top}=\left( \mathbf{AQ} \right)\left( \mathbf{AQ} \right)^{\top}=\mathbf{AQ}\mathbf{Q}^{\top}\mathbf{A}^{\top}=\mathbf{A}\mathbf{A}^{\top}.$$

Thus pairwise relations among samples in the latent space are unaffected. Likewise, the loading Gram matrix is invariant:

$$\mathbf{B}_{\mathrm{rot}}\mathbf{B}_{\mathrm{rot}}^{\top}=\left( \mathbf{BQ} \right)\left( \mathbf{BQ} \right)^{\top}=\mathbf{BQ}\mathbf{Q}^{\top}\mathbf{B}^{\top}=\mathbf{B}\mathbf{B}^{\top}.$$

Hence the feature–feature similarity structure implied by the loadings is unchanged. If $\boldsymbol{\beta}$ is co-rotated as

$$\boldsymbol{\beta}_{\mathrm{rot}}:=\mathbf{Q}^{\top}\boldsymbol{\beta},$$

then

$$\hat{\mathbf{y}}=\boldsymbol{A\beta}=\left( \mathbf{AQ} \right)\left( \mathbf{Q}^{\top}\boldsymbol{\beta} \right)=\mathbf{A}_{\mathrm{rot}}\boldsymbol{\beta}_{\mathrm{rot}},$$

so fitted values for $\mathbf{y}$ are also unchanged.

More generally, right-orthogonal rotation acts within the $R$-dimensional component space of each loading row, preserving its Euclidean norm:

$$\parallel\mathbf{B}_{j,:}\parallel_{2}=\parallel\left( \mathbf{BQ} \right)_{j,:}\parallel_{2}=\parallel\left( \mathbf{B}_{\mathrm{rot}} \right)_{j,:}\parallel_{2} \text{for all }j\in\{1,\ldots,J\}.$$

Consequently, feature summaries based on row norms (and likewise blockwise Frobenius norms) are basis-free.

To see how overlap affects component-wise interpretation, consider an orthogonal re-basing restricted to two components $r\neq s$, written as

$$\mathbf{Q}_{rs}=\left( \begin{matrix} \ddots& & & & \\ & q_{11} & q_{12} & & \\ & q_{21} & q_{22} & & \\ & & & \ddots& \end{matrix} \right), \text{with }\left( \begin{matrix} q_{11} & q_{12} \\ q_{21} & q_{22} \end{matrix} \right)^{\top}\left( \begin{matrix} q_{11} & q_{12} \\ q_{21} & q_{22} \end{matrix} \right)=\mathbf{I}_{2}.$$

Then, for each feature $j$,

$$\left( \begin{matrix} b_{jr}^{\mathrm{rot}} \\ b_{js}^{\mathrm{rot}} \end{matrix} \right)=\left( \begin{matrix} q_{11} & q_{12} \\ q_{21} & q_{22} \end{matrix} \right)\left( \begin{matrix} b_{jr} \\ b_{js} \end{matrix} \right),$$

that is,

$$b_{jr}^{\mathrm{rot}}=q_{11}b_{jr}+q_{12}b_{js}, b_{js}^{\mathrm{rot}}=q_{21}b_{jr}+q_{22}b_{js}.$$

Therefore

$$(b_{jr}^{\mathrm{rot}})^{2}+(b_{js}^{\mathrm{rot}})^{2}=\left( q_{11}b_{jr}+q_{12}b_{js} \right)^{2}+\left( q_{21}b_{jr}+q_{22}b_{js} \right)^{2},$$

which expands to

$$\left( q_{11}^{2}+q_{21}^{2} \right)b_{jr}^{2}+2\left( q_{11}q_{12}+q_{21}q_{22} \right)b_{jr}b_{js}+\left( q_{12}^{2}+q_{22}^{2} \right)b_{js}^{2}.$$

Since the $2\times2$ block is orthogonal,

$$q_{11}^{2}+q_{21}^{2}=1, q_{12}^{2}+q_{22}^{2}=1, q_{11}q_{12}+q_{21}q_{22}=0,$$

and hence

$$(b_{jr}^{\mathrm{rot}})^{2}+(b_{js}^{\mathrm{rot}})^{2}=b_{jr}^{2}+b_{js}^{2}.$$

Thus, orthogonal re-basing can redistribute a feature’s contribution across components without changing its total contribution to the corresponding latent subspace. The individual signed loadings need not be preserved: orthogonal re-basing may alter both their magnitudes and their signs, even though the combined within-subspace contribution remains unchanged.

This has an important consequence for interpretation. Although the reconstruction $\mathbf{A}\mathbf{B}^{\top}$, the sample–sample relations $\mathbf{A}\mathbf{A}^{\top}$, the feature–feature relations $\mathbf{B}\mathbf{B}^{\top}$, and the fitted values $\hat{\mathbf{y}}=\boldsymbol{A\beta}$ (under co-rotation of $\boldsymbol{\beta}$) remain invariant, the sparsity pattern of individual columns of $\mathbf{B}$ is not necessarily conserved. In particular, orthogonal re-basing can redistribute loadings across components, so exact component-wise support patterns may change even when the broader multiblock signal is unaffected. Overlap in the loadings therefore increases the number of observationally equivalent component-wise representations.

When the CLD constraints do not fully determine a unique basis, this allows for rotations and exact component-level interpretation becomes less well defined, even though the broader conclusions about shared signal, cross-block structure, and predictive behaviour are unaffected. In this case, a post-hoc orthogonal rotation such as varimax may be viewed as an interpretability convention: it selects one basis within the same identified latent subspace, without changing the fitted subspace itself or the associated global conclusions.

**4. Exponential family machinery of PESCA is still intact**

The loss functions presented in the main text are simplified versions of those described in Song et al. (2020), included to aid interpretation. Nevertheless, the full estimation machinery for alternative exponential-family distributions, including Bernoulli and Poisson, remains available. As a demonstration, we binarised the ASV block of the Abedini dataset and applied PESCAR. The model converged and produced conclusions broadly consistent with those obtained under the Gaussian-only analysis. This should be interpreted with some caution, however, since binarisation of a count or quantitative matrix inevitably results in substantial information loss. The apparent stronger fit of the binary block reflects the relative ease of modelling this simplified structure.

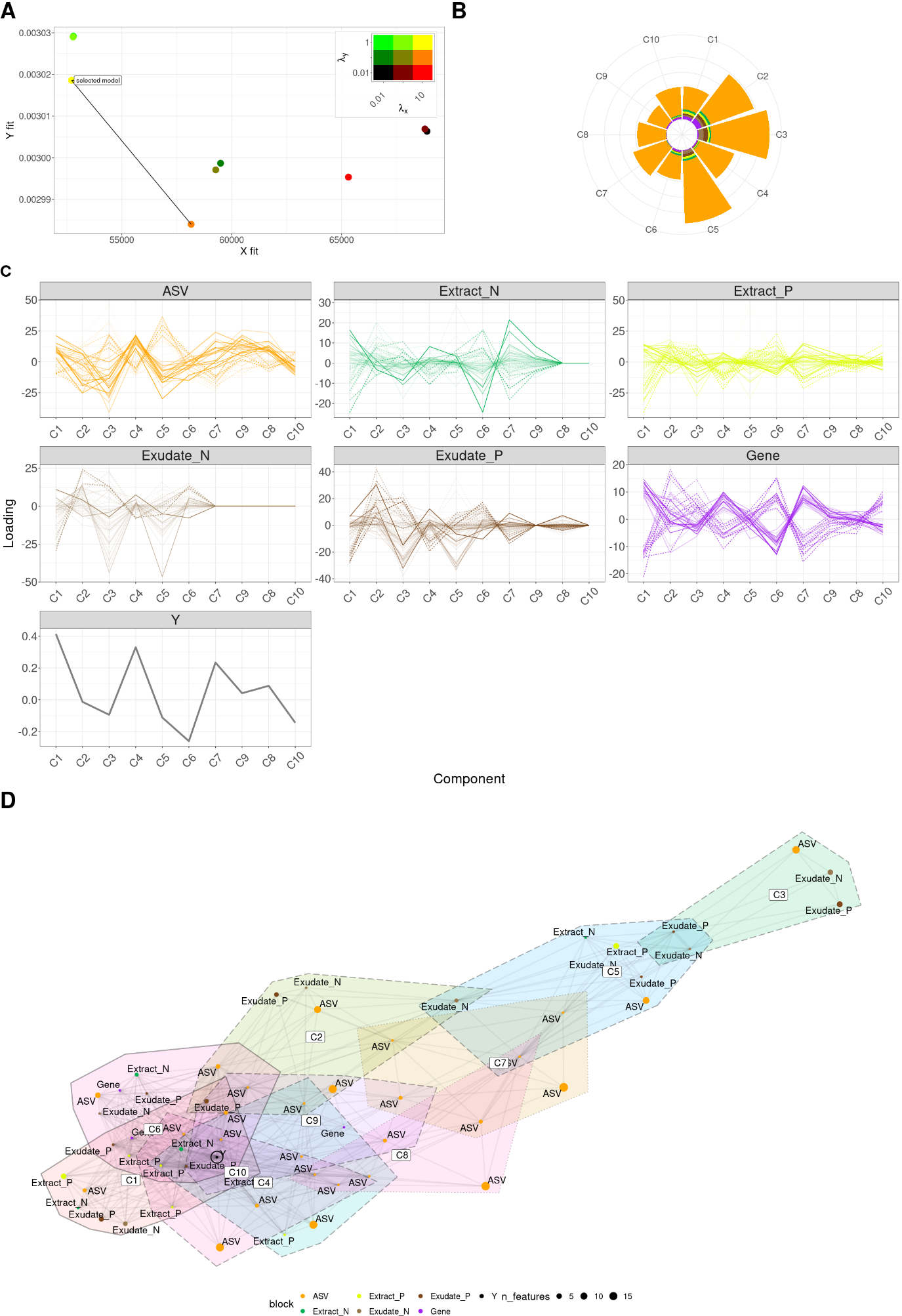

***Supplementary Figure 2:*** demonstration of PESCAR on real data with binarised ASV block. **A**; Model selection based on the Pareto front over the $\lambda_{x}$and $\lambda_{y}$grid search, indicated in plot inset; redness corresponds to $\lambda_{x}$, greenness corresponds to $\lambda_{y}$, yellow indicates both penalties in use. **B**; Radar plot showing the variance explained by each component in each block, used to identify common, local, and distinct component structure. **C**; Block-wise loading profiles of the selected model across all components, showing the feature loading patterns within each block, line opacity scaled by **y**-relatedness (f). **D**; Hypergraph representation of the full model derived from overlap in the nonzero loading support across components, summarising the higher-level topology of the fitted model, node size indicates number of features represented, edge width indicates the number of components that a pair of nodes are both nonzero.
